# Direct Crystal Formation from Micronized Bone and Lactic Acid: The Writing on the Wall for Calcium-Containing Crystal Pathogenesis in Osteoarthritis?

**DOI:** 10.1101/384669

**Authors:** Anna A. Bulysheva, Nardos Sori, Michael P. Francis

## Abstract

**Introduction:** Pathological calcium-containing crystals accumulating in the joints, synovial fluid, and soft tissues are noted in most elderly patients, yet arthritic crystal formation remains idiopathic. Interestingly, elevated lactic acid and bone erosion are frequently among the comorbidities and clinical features of patients with highest incidence of crystal arthropathies. This work shows that bone particulates (modeling bone erosion) dissolve in lactic acid and directly generate crystals, possibly presenting a mechanism for crystal accumulation in osteoarthritis.

**Methods and Results:** Micronized human bone (average particle size of 160μm x 79μm) completely dissolved in lactic acid in 48 hours, and in synovial fluid with 500mM lactic acid in 5 days, generating birefringent rhomboid and rod-shaped crystals. SEM analysis with energy dispersive x-ray spectroscopy of these crystals showed average dimensions of around 2μm x 40μm, which contained oxygen, calcium and phosphorous at 8.64:1.85:1. Raman spectroscopy of the generated crystals further showed 910/cm and 1049/cm peaks, aligning with calcium oxalate monohydrate and calcium pyrophosphate, respectively.

**Conclusions:** This work shows that lactic acid and micronized mineralized bone together directly generate calcium-containing crystals. These observations may provide insights into the elusive etiology of arthritis with crystal involvement, possibly indicating lactic acid as a clinical target for treatment.

## 1. Introduction

Osteoarthritis (OA) is the most prevalent human joint disorder and a leading cause of disability^1^. However, with its complex and likely multifactorial pathogenesis, OA remains idiopathic. Poor understanding of OA etiology has meant poor therapeutic options^2^, with progressive destruction leading to total joint replacement. At least 70% of OA synovial fluid samples have calcium-containing crystals^3^, yet whether crystals are at the root of OA or are downstream in relation to joint damage is under debate^4^. Pathological calcium and phosphate containing crystals in osteochondral regions of joints, in the surround joint soft tissue and in synovial fluid are commonly noted in most forms of osteoarthritis^5,6^. Basic calcium phosphate (BCP) and calcium pyrophosphate dehydrate (CPP) are the most commonly found crystals in articular cartilage and synovial fluid^7^, along with, very rarely, calcium oxalate monohydrate (COM)^8^. However, multiple species of crystals are often found together in the same patient sample^8^.

Besides gout, calcium pyrophosphate deposition disease (CPPD) (or pseudogout) is the most common idiopathic crystal arthropathy^5,6^, with knee as the most commonly involved joint, yet with polyarticular crystal deposition seen in two-thirds of patients. CPPD is estimated to affect 15% of people aged 65-74, and 50% of people age 84 or older^5,9^. CPP crystals present as rhomboid or rod-shaped structures and are weakly *positive* in birefringence under compensated polarized light microscopy, while the crystals are *non-birefringnent* for BCP deposition diseases^10^. There is no known effective method of removing calcium-containing crystals deposited in the synovium, bound to cartilage (common in CPP) or soft tissues.

The non-apatitic carbonate and phosphate groups on bone minerals (calcium phosphate [Ca_3_(PO_4_)_2_] or hydroxyapatite [Ca_10_(PO_4_)_6_(OH)_2_]) are reported to be very biologically active, which may lead to dissolution of the hard tissue such as in normal remodelling^11^. CPP (Ca_2_PO_7_) or BCP may thus be generated as crystals from the breakdown of bone in the presence of reactive lactic acid (C_3_O_6_H_3_) or lactate (CH_3_CH(OH)COO^-^). Further, CPPD is reportedly somewhat difficult to distinguish from erosive OA clinically ^12^, linking bone particulate to arthritic crystals.

In humans, normal resting lactic acid levels range from 0.5-1.8 mmol/L. However, when CPP crystals are detected, 4.2-11.2mmol/L of lactic acid has been reported, with elevated pH^13^, and up to 10-24 times normal levels in the synovial fluid for rheumatoid arthritis and OA^13^. Lactic acid elevation is further a recurring clinical feature in crystal arthropathy comorbidities **(Table 1)**, along with bone erosion. It is thus hypothesized that lactic acid, or lactate, and eroded bone (modeled here with human micronized bone particulate) directly generate calcium and phosphate-containing crystals, suggesting joint lactic acidosis is directly mechanistically linked to crystal arthropathies. Supporting this hypothesis, this work reports an original model system of micronized mineralized bone suspended in synovial fluid in the presence of acids to study bone-derived crystal formation.

**Table 1.** Crystal Arthropahty Comorbidities Linked with Lactic Acid and Bone

| **Crystal Arthropathy Comorbidities** | **Major Clinical Manifestation** | **Potential Lactic Acid & Eroded Bone Relationships** |
| --- | --- | --- |
| Gout | Urate crystal formation | Reduced uric acid excretion coincides with increased lactic acid concentration in blood |
| Hemochromatosis | Excess iron storage | Elevated lactic acid consumption enhances iron uptake |
| Hypercalcemia | Excess blood calcium level | Possible eroded bone link |
| Hyperparathyroidism | Overactive parathyroid gland | Increase lactic acid yields increased free ionized calcium, which is a diagnostic for pHPT |
| Hypocalciuric Hypercalcaemia | Elevated Serum Calcium (Above 10.2 mg/dl) | Increased bone resorption |
| Hypomagnesemia | Low Blood Magnesium Level (<0.7mmol/L) | Associated with metabolic lactic acidosis |
| Hypophosphatasia | Skeletal hypomineralization or osteomalacia | Elevated endogenous inorganic pyrophosphate noted |
| Hypothyroidism | Underactive thyroid gland | Excess lactic acid is produced even at rest |
| Infection | Sepsis from microbes | Lactic acid accumulation |
| Ischemic Heart Disease | Loss of blood flow and necrosis of myocardium | Lack of oxygen shifts myocardium to anaerobic metabolism, elevating lactic acid |
| Osteoarthritis | Degenerating cartilage and subchondral bone | Eroded bone present micronized bone particles |
| Rheumatoid Arthritis | Joint lining inflammation and bone erosion | Lactic Acid builds up to repair inflamed muscles and joints, and erosion + fragments of bone noted |
| Surgery | Soft tissue ablation | Lactic acid is the main component of Lactated Ringers & Hartmann's solution |
| Trauma | Tissue injury | Lactic acid levels strongly associated with morbidity in level 1 trauma centers |

## 2. Methods

### 2.1 Bone Particulate Production

Human bones from three donors with informed research consent (generous gift via the NDRI) were debrided, with the marrow and trabecular bone removed. Washed bone was micronized in a TekMar^®^ grinder and sieved to <125μm and 125-750μm. Portions of the bone from the two sieved size ranges were demineralized using 0.1M hydrochloric acid (SigmaAldrich, St. Louis, MO) and rinsed with phosphate buffer to neutralize (SigmaAldrich, St. Louis, MO).

### 2.2 Synovial Fluid Collection and Crystal Formation

Arthrocentesis was performed postmortem on the knees joint of 4 recently deceased male donors (70-80 years old) with research consent and assessed microscopically for crystals (n=1 with crystals, n=3 without) by a pathologist. Synovial fluid was stained with alizarin red S (SigmaAldrich, St. Louis, MO) to ensure absence of hydroxyapatite, as compared to control samples. Normal synovial fluid was combined with 0, 10, 50, 100 or 500 mM of lactic acid, and with 100mg/ml of micronized mineralized bone. These samples were incubated at 37°C with and without shaking at 1000RPM shaking (Eppendorf Thermomixer R, Hamburg, Germany) then observed microscopically with normal and polarizing light daily for two weeks.

### 2.3 Acid Treatments of Bone and Minerals

100mg/ml of mineralized and demineralized bone particulate <125μm and 125-750μm from 3 donors were stirred at 750RPM combined with either 1M or stock lactic acid, phosphoric acid, oxalic, hydrochloric acid, acetic acid, sulfuric acid, formic acid, citric acid, or trifluoroacetic acid (all from SigmaAldrich, St. Louis, MO) at either 4°C, room temperature (~21°C), or 37°C. Samples were taken a 2hrs, 8hrs, 16hrs, and every 24hrs for 2 weeks for microscopic evaluation. The solubility of BCP, HA, and CPP (Sigma-Aldrich, St. Louis, MO) was also tested in each acid at 100mg/ml and imaged under normal and polarized light. Samples were examined for the appearance of crystals by an Olympus BX41 microscope with a polarizer. CPPD crystals from arthrocentesis were imaged as positive controls.

### 2.4 Scanning Electron Microscopy (SEM)

SEM analysis was performed on a JEOL JSM-6060LV at Jefferson Labs (Newport News, VA). Crystals generated from lactic acid dissolved mineralized bone were vacuum filtered (Whatman #4) and dried under a fume hood for 24 hours to partially dry the samples. Mineralized dry bone particulates were sputter coated with gold for SEM imaging. Samples for EDS (uncoated) were imaged at low pressure (100Pa).

### 2.5 Quantitative SEM Image Analysis

ImageJ64 (NIH Shareware) was used to measure the average bone particle sizes and average crystal dimensions. The averages are represented as the mean ± the standard deviation.

### 2.6 Raman Spectroscopy

Mineralized bone particulate and lactic acid dissolved bone particulate were analyzed by Raman spectroscopy on a Thermo Scientific smartRaman 532 nm with OMNIC software at Virginia Commonwealth University (Richmond, VA). Crystals of calcium pyrophosphate, hydroxyl apatite, calcium phosphate (Sigma-Aldrich) were used dry and at 100mg/ml in lactic acid to generate a Raman reference library to compare the crystals generated from bone in lactic acid sample. Lactic acid alone was analyzed to assess background interference. Micro-Raman at 633nm was performed at ChemXLerate (Ann Arbor, MI) for bone in lactic acid samples.

## 3. Results

### 3.1 Crystal generation from bone in lactic acid

Of the nine acids tested, only lactic acid was found to dissolve and liquefy mineralized (but not demineralized) bone particulate in the sub 125-micron diameter particle size range (henceforth referred to as “bone” per this proposed modeled system of eroded bone mineral interacting with elevated local or systemic lactic acid). Eroded bone treated with lactic acid samples presented as turbid, milky white appearance, similar to clinical samples with crystals. Lactic acid treated bone formed rhomboid crystals in approximately 16 hours at 37°C, forming rod-shaped crystals in around 30 hours at room temperature, and forming rhomboid crystals around 48 hours at 4°C, from all 3 donors. Phosphoric and trifluoroacetic acids dissolved mineralized and demineralized bone but did not form crystals.

Under normal and polarized light microscopy, a donor knee synovial fluid sample with pathologic crystals was found containing positively birefringent rod-shaped crystals **(Figure 1A,B)** as a reference control. Adding 500mMol of lactic acid with 100mg/mL of bone to the healthy synovial fluid (negative control, no crystals present initially) generated birefringent rod-shaped crystals in the synovial fluid by 5 days of 37°C incubation **(Figure 1C,D)**, with groups exposed to a lower concentration of lactic acid (100mM) and mineralized bone in normal synovial fluid formed rod-shaped crystals at 2 weeks (not shown). The dissolved rhomboid and rod-shaped crystals from bone exhibited positive birefringence in both 500mMol lactic acid in human synovial fluid **(Figure 1D)**, and also formed crystals with micronized bone directly dissolved into 85% lactic acid **(Figure 1E,F)**, comparable to control diseased synovial fluid with crystals **(Figure 1B)**. Birefringence was observed for most, but not all, crystals present. Titration of deionized water to the crystal samples progressively resulted in shrinkage and dissolution of the crystals.

**Figure 1.**
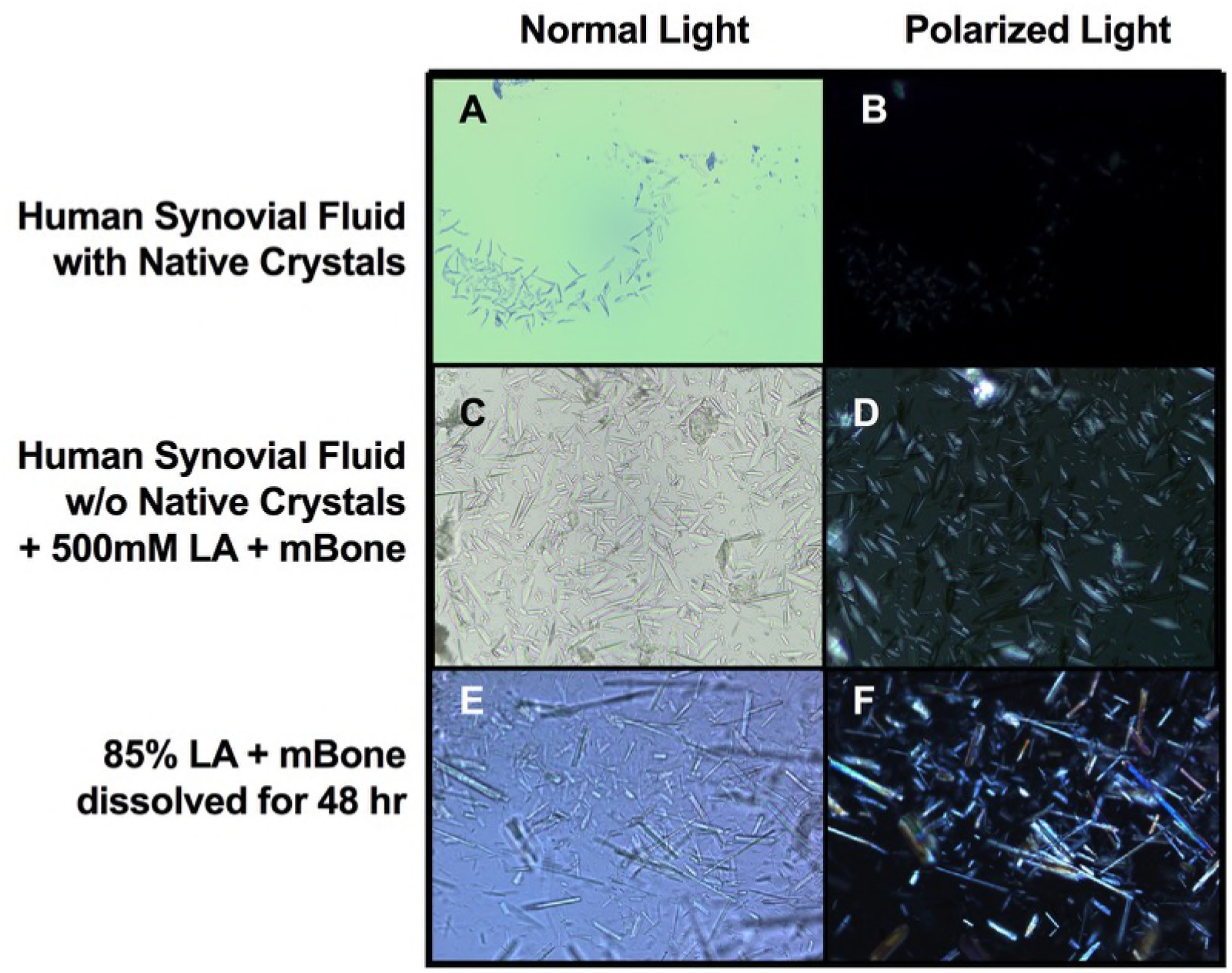
Crystals generated from lactic acid treated micronized bone. Human knee synovial fluid was collected from a donor that was diagnosed with CPPD, containing crystals seen under normal **(A)** and polarized light **(B)** microscopy. Synovial fluid from a second male donor without crystals present was mixed with 500mMol of lactic acid (LA) and 100mg/ml of micronized mineralized bone (mBone), which produce birefringent crystals after 5 days of incubation at 37°C **(C, D)**. Using 85% lactic acid exposure to micronized mineralized bone, rhomboid crystals are noted to form initially (around 18 hours) followed by rod shaped crystals by 48 hours in solution **(E)**. These crystals in lactic acid also exhibit birefringence under polarized light **(F)**, and are further stable in this form for at least a month.

Lactic acid dissolved bone at 100mg/ml from two unique donors imaged by SEM showed relatively homogeneous crystals with average length of 39.53μm by 2.07μm width, relative to the starting bone particulate averaging 160.36μm x 78.79μm (**Figure 2A-C**).

**Figure 2.**
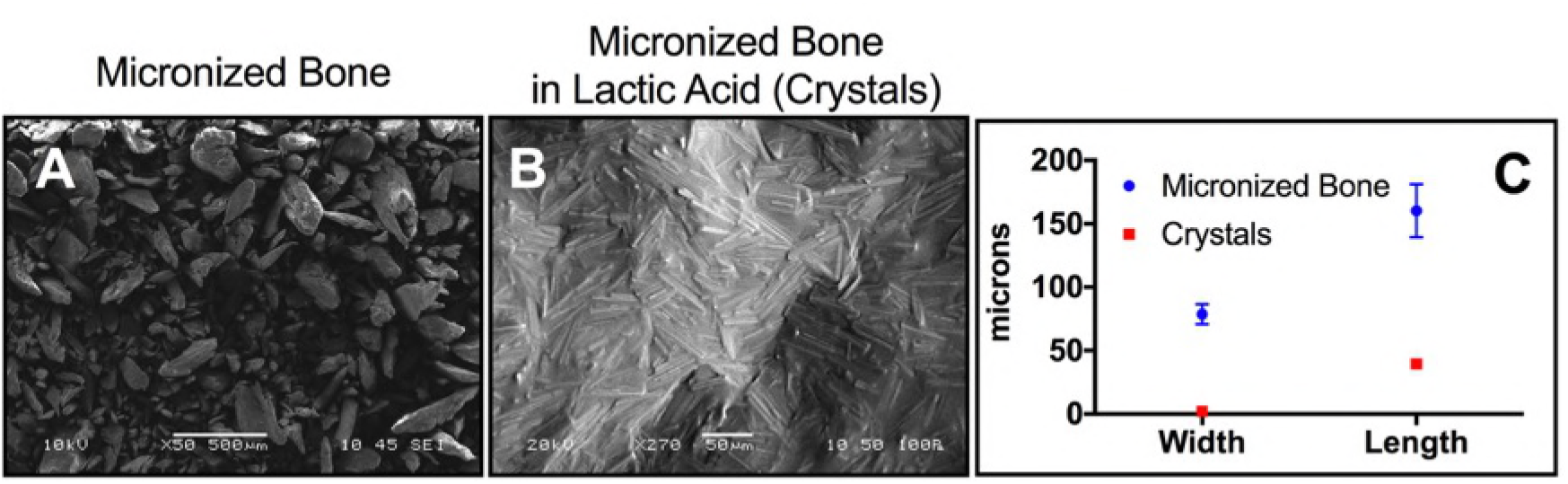
SEM imaging of bone particulates and lactic acid generated crystals from mineralized bone. Representative images of micronized mineralized human bone **(A)**, and the crystals generated from lactic acid exposure of micronized mineralized bone (still partially wet in acid) **(B)** are shown by SEM. The bone particle and resulting crystal size distribution is further quantified **(C)**.

### 3.2 Crystal Elemental Analysis

SEM analysis with EDS of micronized mineralized bone alone **(Figure 3A)** is compared to crystals generated from bone in synovial fluid with 500mMol lactic acid **(Figure 3B)**, the latter of which showed atomic percentages of calcium and phosphate in the crystals at ratio of O:Ca:P at 8.64:1.85:1 (n=5). Micro-Raman spectroscopy (633nm) of crystals from bone in lactic acid did not detect the peak for hydroxyapatite at 960/cm yet the spectrum does contain the dominant 1049/cm band used to diagnose CPPD clinically **(Figure 3C)**, suggesting the presence of Ca-O and P-Ca bonds. This 1049/cm peak was also seen in CPP crystals controls, but not BCP in lactic acid, as tested by Raman spectroscopy at 532nm (not shown).

**Figure 3.**
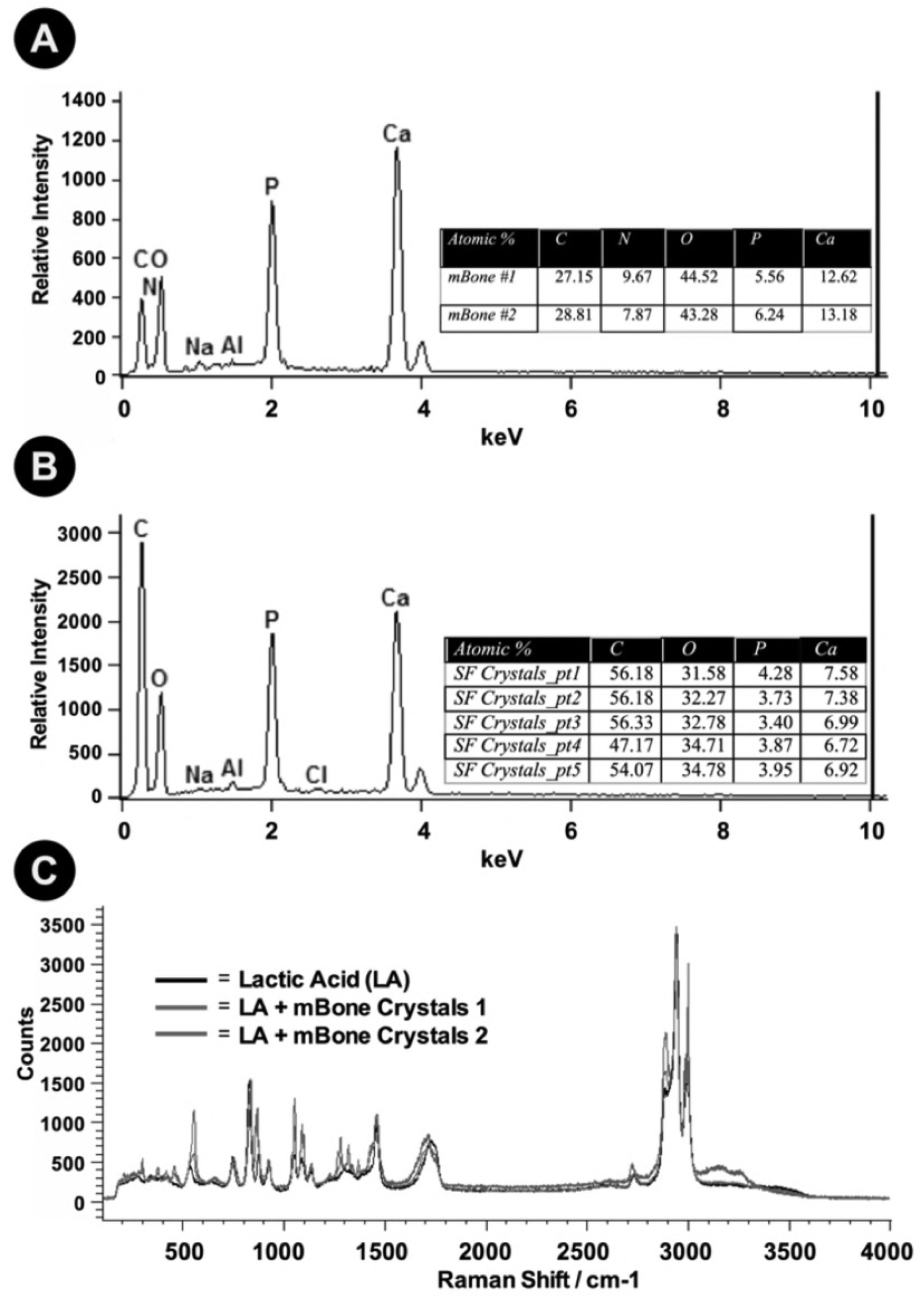
Raman and energy-dispersive X-ray spectroscopy analysis of crystals generated from micronized bone in lactic acid. Representative EDS spectra and quantitative results of the untreated micronized mineralized bone particulate **(A)** and lactic acid treated crystals **(B)** are shown. Micronized mineralized human bone (100mg/ml) dissolved in synovial fluid with 500mMol of lactic acid was analyzed by EDS after vacuum drying, which, while still slightly wet with lactic acid, presented an atomic percentage for calcium, phosphate, and oxygen between that of BCP and CPP. EDS was performed on two individually processed donors for each condition (n=2), yielding elemental composition results that were not significantly different. Micro-Raman spectroscopy analysis was performed on lactic acid (LA) with micronized mineralized bone-generated crystals (labeled as “LA + mBone Crystals 1 & 2”) and compared to lactic acid alone, with imaging targeted at a single crystal for analysis. A dominant peak seen at 1049/cm (using 633nm spectroscopy) is shown in the experimental traces **(C)**, which is a peak commonly reported for calcium pyrophosphate crystals clinically. Further, the hydroxyapatite peak (commonly reported at 960/cm) is absent, yet a slight peak is noted around 900/cm, indicative of calcium oxalate monohydrate.

### 4. Discussion

Better understanding of the causes of crystal formations are still needed to reduce, treat, or prevent the formation and deposition of arthritic crystals. Lactic acid has been considered a byproduct or correlated feature of arthritis involving crystals for decades^13^. This work presents a hypothetical pathogenesis for calcium-containing crystal, where micronized bone (modeling eroded bone) and lactic acid are directly shown to generate crystals analogous to those common in various forms of OA **(Figure 4)**, being the first time this has been experimentally reported.

**Figure 4.**
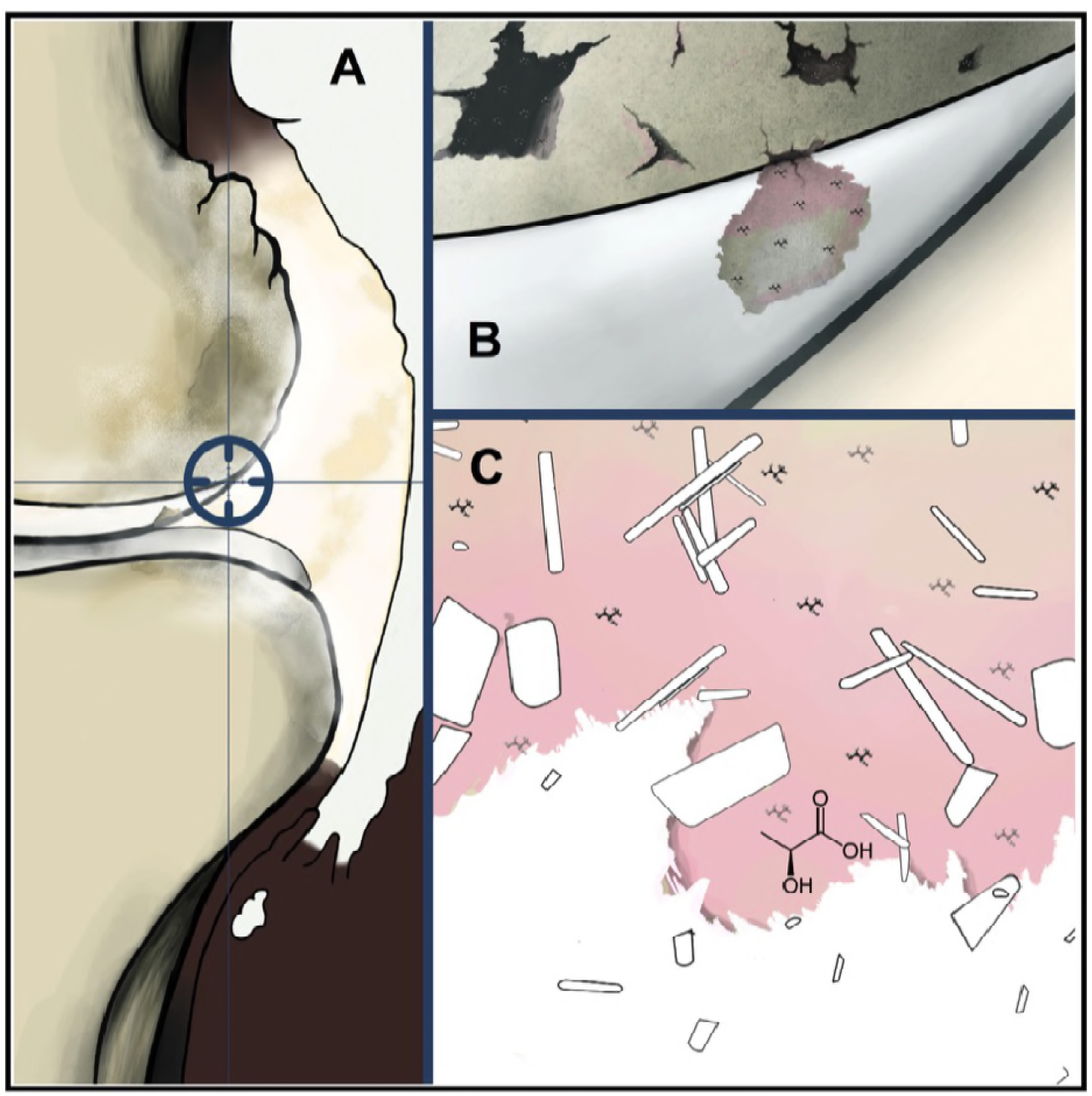
Proposed Mechanism of Arthritic Calcium-Containing Crystal Formation in Joints. **(A)** Illustration showing the theoretical pathological origin of calcium-containing crystals as generated in the synovial space of the knee from the interaction of eroded bone (B) and lactic acid (C) resulting in progressive degradation of the injured bone to generate calcium-containing crystals.

Bone is solubilized in lactic acid to form crystals, as may be occurring in arthritic bone and cartilage from erosions. For synovial fluid, 500mMol of lactic acid was required in around 5 days of exposure to generate rhomboid and rod-like crystals **(Figure 1)**. Below, we propose a possible mechanism for the reaction of lactic acid and hydroxyapatite **(Scheme 1)** to form crystals. In the first step, lactic acid reacts with hydroxyapatite at the site of calcium forming intermediate 1. The dissociation of intermediate 1 forms a precursor of the final product with the formation of intermediate 2. In the final state, the presence of oxygen in the surrounding interacts with intermediate 2 to form calcium phosphate and calcium oxalate monohydrate.

**Scheme 1.**
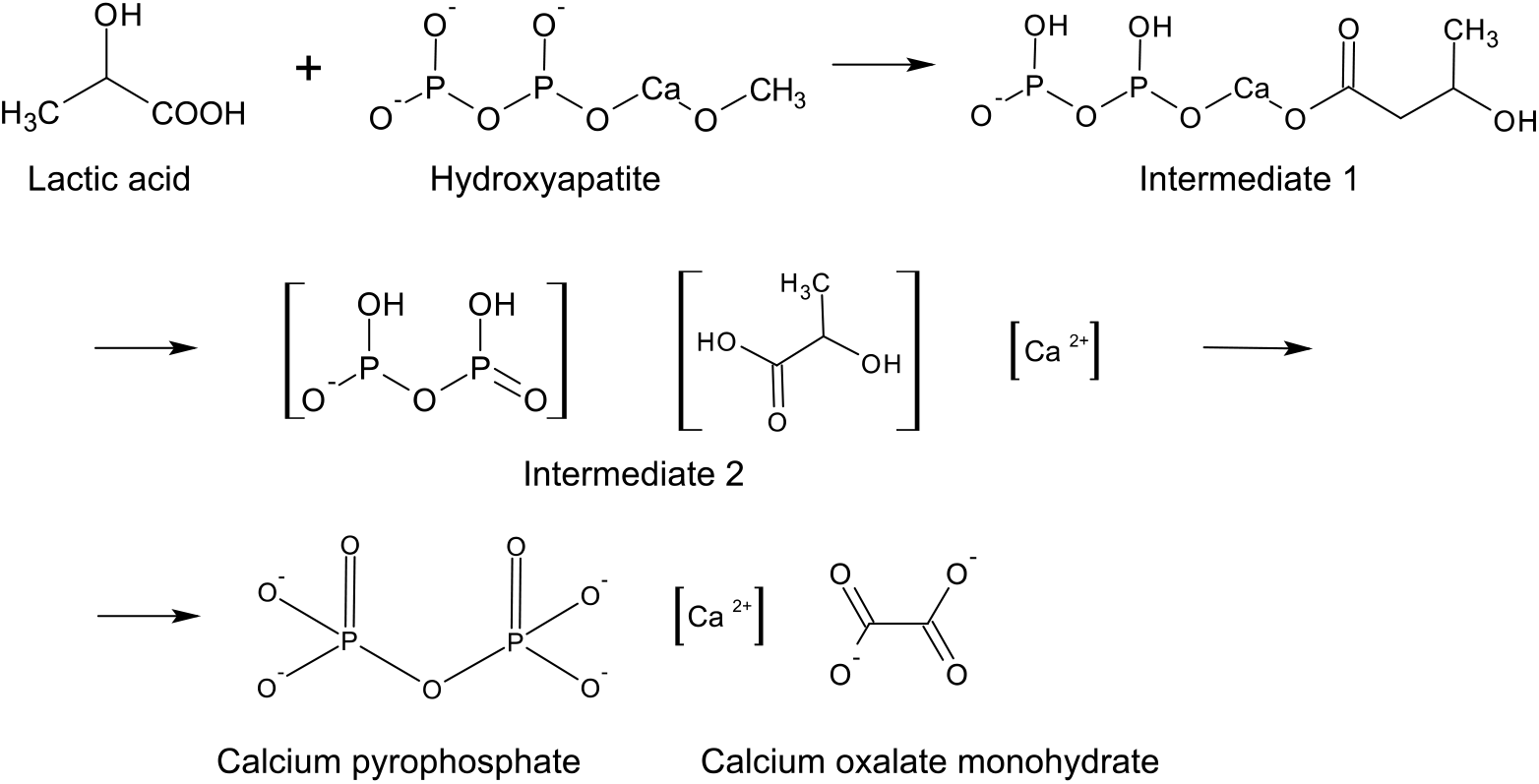
Proposed mechanism of crystal formation. Synthesis of calcium pyrophosphate and calcium oxalate monohydrate from lactic acid and hydroxyapatite.

In this study, while the lactic acid concentrations used are higher than physiologic, the reaction kinetics of the acid with bone would occur at lower concentrations as well, albeit with requisite additional reaction time. The higher concentrations of lactic acid tested where thusly chosen to speed the chemical reaction to facilitate analyses. Lactic acid dissolved bone crystals dissolved under increased hydration, similar to clinical reports^13^ suggesting hydration as a possible therapeutic consideration and perhaps related to the 33% decrease in body water content in the elderly compared to the young^14^. While the size of the crystals generated in this study are larger than those seen clinically for CPP or BCP, the high saturation of bone and higher acid concentration tested may allow for crystallization to occur beyond what is common clinically.

SEM-EDS results of micronized bone in 85% lactic acid and of micronized bone in 500mMol of lactic acid in synovial fluid show the presence of oxygen:calcium;phosphate in the crystals at a ratio of 8.64:1.85:1 **(Figure 3)**. This ratio is not perfectly fitting with CPP (Ca:P at 1:1) or BCP (Ca:P at 2:1), yet as the samples are of dissolved bone crystals in wet solution, free bone salts in the solution may be interfering with the measurements. Raman spectroscopy was thus used to further identify the crystals, where the spectra indicated a dominant peak at 1049/cm, positively identifying CPP, whereas the hydroxyapatite peak at 960/cm is absent, with a slight peak at around 910/cm suggesting calcium oxalate crystals **(Figure 3)**. The reference library generated from all Raman samples at 532nm suggested that lactic acid dissolved bone samples were most similar to CPP controls.

Review of the literature **(Table 1)** revealed elevated lactic acid levels, bone resorption and erosion frequently co-associated with crystal arthropathy comorbidities, further linking lactic acid and crystal formation. Subsequent epidemiological studies assessing lactic acid involvement in arthritis are clearly warranted. While treating cartilage and bone erosion in arthritis has proven incredibly challenging, approaches addressing lactic acid reduction may hold promise. Care strategies to reduce lactic acid levels to potentially inhibit or reverse crystal formation may include improved hydration, reduced lactic acid-containing food consumptions, increased physical activity, dietary supplementation (e.g. beta alanine, taurine) and possible pharmacological interventions.

For additional future directions, it may prove informative to test synovial membrane cells from normal and OA individuals to assess if either tends to produce more lactic acid. Interestingly, synovial fluid further contains the lactic acid dehydrogenase (LDH) enzyme, suggesting importance of joint-lactic acid balance^15^. LDH therapy (e.g. gene therapy) may further be tested in animal models for controlling crystal-containing OA. Further, lactic acid may be administered to animals to potentially induce bone erosion, OA and crystal formation, possibly even spurring rheumatoid arthritis symptoms.

## 5. Conclusion

We report on a proposed model for the etiology of osteoarthritis with crystal involvement, where eroded bone particulate in the presence of lactic acid are shown to directly generate calcium-containing crystals. These observations are significant because of the current idiopathic nature of calcium-containing crystal arthropathies. This work shows that lactic acid, a common acid in the body that accumulates with age and with disease, is remarkably found to experimentally dissolve bone particulate and he dissolved bone salts form stable crystals that resemble those found in some forms of osteoarthritis. These results and the hypothesis herein present a plausible direct cause of osteoarthritis with crystal involvement from local eroded bone and the elevated lactic acid in the synovial fluid and systemically, as is commonly reported with osteoarthritis.

